# LDLR-mediated targeting and productive uptake of siRNA-peptide ligand conjugates *in vitro* and *in vivo*

**DOI:** 10.1101/2023.02.03.526778

**Authors:** Broc B., Varini K., Sonnette R., Pecqueux B., Benoist F., Thomas M., Masse M., Mechioukhi Y., Ferracci G., David M., Temsamani J., Khrestchatisky M., Jacquot G., Lécorché P.

## Abstract

siRNAs have become one of the most promising therapeutic agents because of their specificity and their potential to modulate the expression of gene-related diseases. Any gene of interest can be potentially up or down-regulated, making RNA-based technology the healthcare breakthrough of our era. However, the functional and specific delivery of siRNAs into tissues of interest and into the cytosol of target cells remains highly challenging, mainly due to the lack of efficient and selective delivery systems. Among the variety of carriers for siRNA delivery, peptides have become essential candidates because of their high selectivity, stability and conjugation versatility. Here, we describe the development of molecules encompassing siRNAs against *SOD1*, conjugated to peptides that target the LDLR, and their biological evaluation both *in vitro* and *in vivo*.

GRAPHICAL ABSTRACT

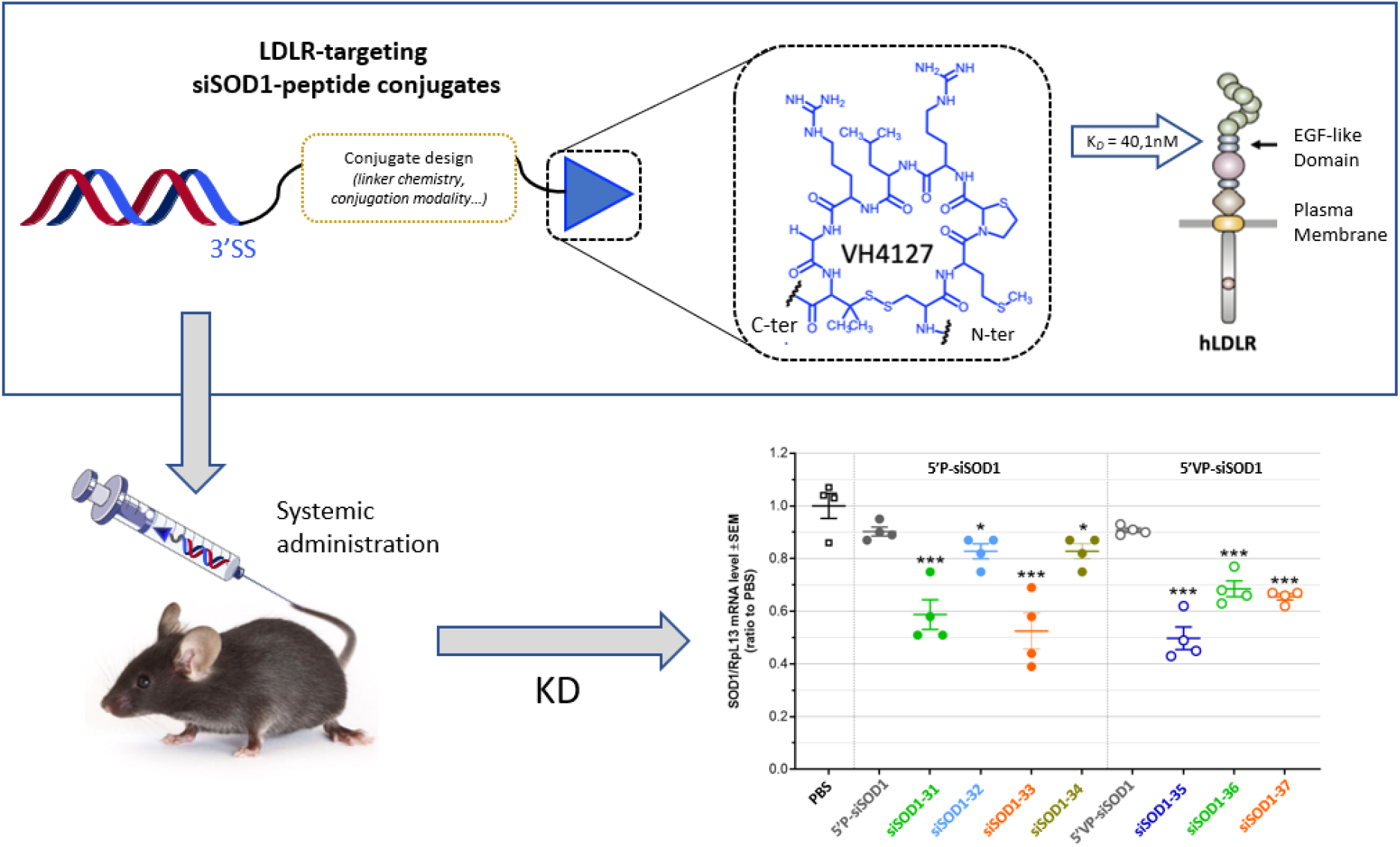

## INTRODUCTION

RNA interference (RNAi) has become a robust tool to silence, in a highly specific manner, genes of interest in mammalian cells. The underlying mechanism of RNAi is based on the uptake by the cytoplasmic RNA-induced silencing complex (RISC) of a small (21-24 nucleotides in length) interfering RNA (siRNA) that binds in a specific base-pairing manner with its complementary mRNA sequence, thereby triggering its degradation and suppressing expression of a disease-causing protein^1–3^. Theoretically, therapeutic siRNAs have the potential to treat any gene-related pathology thanks to their potency, specificity^5^, and duration of effect^6^.

However, because the small size of unconjugated stabilized siRNAs is within the limit of active filtration, they spontaneously present a preferential tropism for excretory organs when systemically administered^7,8^ and are expected to be cleared in majority (~85%) through the kidney in a short period of time. The specific delivery of siRNAs in tissues of interest remains a scientific lock, mainly due to the lack of efficient and selective delivery systems^9,10^. From lipid and polymer nanoparticles^11–14^, that encapsulate free siRNA, to well-defined and stable molecular Ligand-siRNA conjugates, a wide variety of delivery systems are currently on the study bench to fully unlock the potential of siRNAs.

While numerous cell surface markers can be targeted to allow tissue-specific enrichment of carrier-drug conjugates, only a few examples of ligand/receptor pairs were shown to support efficient and tissue-selective functional delivery of siRNAs^18^. Following the clinical success of GalNAc-siRNA conjugates, that efficiently and specifically target liver hepatocyte ASGPRs allowing potent functional uptake and gene knock-down^19,20^, other receptors supporting constitutive and/or ligand-mediated endocytosis represent attractive cell-surface receptors for siRNAs to reach intracellular compartments at extrahepatic sites^21^. Among them, the LDL receptor (LDLR) is an attractive cell surface target. First, it is endowed with high recycling activity leading to high uptake potential of circulating ligands, as evidenced by its fundamental role in LDL-cholesterol plasma clearance^22^. The underlying subcellular mechanisms have been well described and rely on the efficient endosomal release of LDL in the mildly acidic environment (pH 6.0) of early sorting endosomes (SE)^23–25^. Because the level of LDLR and its uptake capacity correlate with the need for LDL-derived cholesterol in major biological processes, the LDLR displays some degree of tissue specificity that may be exploited in pathophysiological conditions including cancer.

Given the large therapeutic opportunities of LDLR as a relevant cell surface receptor in targeted drug delivery approaches, we identified and optimized a family of peptide-based vectors that target the LDLR and that meet the following requirements: i) high selectivity and nanomolar affinity for both the rodent and human LDLR to allow preclinical studies while managing risk for further clinical studies, ii) minimal sized 8 amino acid peptides, cyclic and chemically optimized for increased stability, iii) absence of competition with the binding of LDL, iv) proven conjugation versatility while retaining LDLR uptake capacity, allowing targeting of a variety of cargos ranging from small organic molecules, peptides, siRNAs and proteins, including antibodies, and v) *in vivo* validation of their specific LDLR-dependent tissue distribution in wild-type or ldlr−/− mice^26,27^.

Among the family of peptide-based vectors we identified, the VH4127 (cyclo[(D)-Cys-Met-Thz-Arg-Leu-Arg-Gly-Pen]) peptide ligand used in this study was described elsewhere^26^. Briefly, it was obtained by high-throughput screening of phage display peptide libraries on engineered cell lines expressing the mouse and human LDLR and was further chemically optimized for optimal LDLR-binding affinity and plasma stability^28^. The VH4127 peptide contains three non-natural amino-acids D-Cys, Thz and Pen, (at position 1, 3 and 8 respectively) and a disulfide bridge between D-Cys and Pen lateral chains (Figure 1-A). LDLR-binding kinetics (on-rate, k_on_, and off-rate, k_off_) of peptide VH4127 assessed using surface plasmon resonance (SPR) allowed determination of its equilibrium dissociation constant (*K_D_*=40,1nM). *In vitro* plasma stability of peptide VH4127 was evaluated by incubation at 2mM in freshly collected mouse blood at 37°C. LC-MS/MS analysis performed on the plasma fractions at the end of indicated time points led to an estimated half-life of 4.27h.

**Figure 1:**
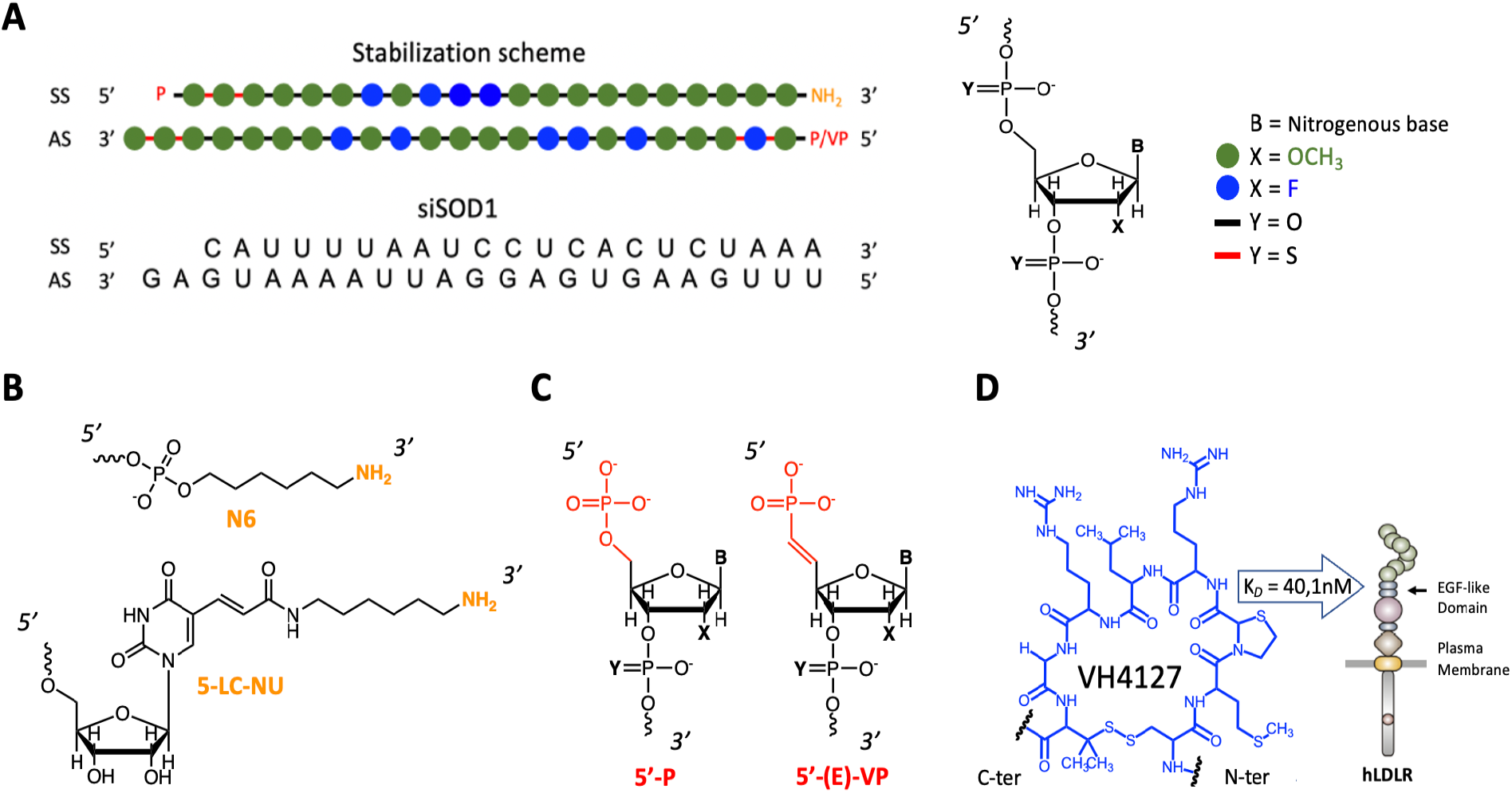
General design of siSOD1-peptide conjugates. **(A)** Detail of the stabilization diagram of siSOD1 duplex. Chemical modifications: green dot = 2’-OMe; blue dot = 2’-F; red line = PS; P = phosphate; VP = vinylphosphonate. **(B)** Detailed structure of the reactive amine at the 3’-end of the sense strand for further conjugation. N6 = 6-carbon aliphatic arm ending with an amine group. 5-LC-NU = 5-Aminohexylacrylamino-Uridine, modified uridine with an aminohexylacrylamine arm at position 5. **(C)** Detailed structure of the 5’-end of the antisens strand. P = standard phosphate; (E)-VP = modified and metabolically stable vinylphosphonate with a double bond in E configuration. **(D)** Detail of the VH4127 peptide sequence (cyclo[(D)-Cys-Met-Thz-Arg-Leu-Arg-Gly-Pen]). Disulfide cyclization occurred between the penicillamine and cysteine lateral chains. VH4127 LDLR-binding affinity: K_D_ = 40,1nM (SPR: Biochip NiHC1000m; mode MCK).

Here, we describe the development of a panel of siRNA-VH4127 peptide conjugates targeting the LDLR and their biological evaluation in both *in vitro* and *in vivo* studies. A siRNA targeting the ubiquitously expressed *SOD1* mRNA was used as a model allowing evaluation of the functional delivery potential of these conjugates.

## RESULTS

### Molecular design and synthesis of siSOD1-peptide conjugates

Design of the siSOD1-peptide conjugates involved three-partners, namely i) the peptide ligand allowing specific tissue-targeting, ii) the pharmacologically active siRNA moiety, and iii) the linker that is essential for linking these two functional entities while retaining their biological functions. Since the VH4127 peptide was previously selected, optimized and validated for its ability to preferentially distribute *in vivo* to LDLR-enriched tissues^26,27^, the present work focused on the siRNA and the chemistry process used for its conjugation to the VH4127 peptide.

We explored different conjugation strategies to evaluate the impact of the different designs on the physico-chemical properties of the conjugates and on their *in vitro* and *in vivo* efficacies. Two different methodologies were investigated: i) an indirect convergent strategy through a strain-promoted azide alkyne cycloaddition (SPAAC) process that requires the first parallel introduction of suitable moieties on both the siSOD1 and the peptide for further click conjugation (Figure 2-A, B) and ii) a direct conjugation by an amide bond. A murine siSOD1 was studied in this work based on the stabilization scheme described by Foster et al.^29^. It encompasses chemical modifications that increase its resistance to nucleases and silencing potency while diminishing potential off-target effects and cytotoxicity. The duplex was composed of a sense strand (SS) of 21 nucleotides, with an hexylamino (N6) modification at the 3’-end, and an antisense strand (AS) of 23 nucleotides with a 2 nucleotide 3’-overhang ending. The N6 modification is a 6-carbon aliphatic arm ending with a reactive primary amine or a 5-LC-NU modification for further chemical functionalization. Finally, both SS and AS sequences contained a 5’-modification, respectively a 5’-phosphate or 5’-vinylphosphonate modification (Figure 1-B).

**Figure 2:**
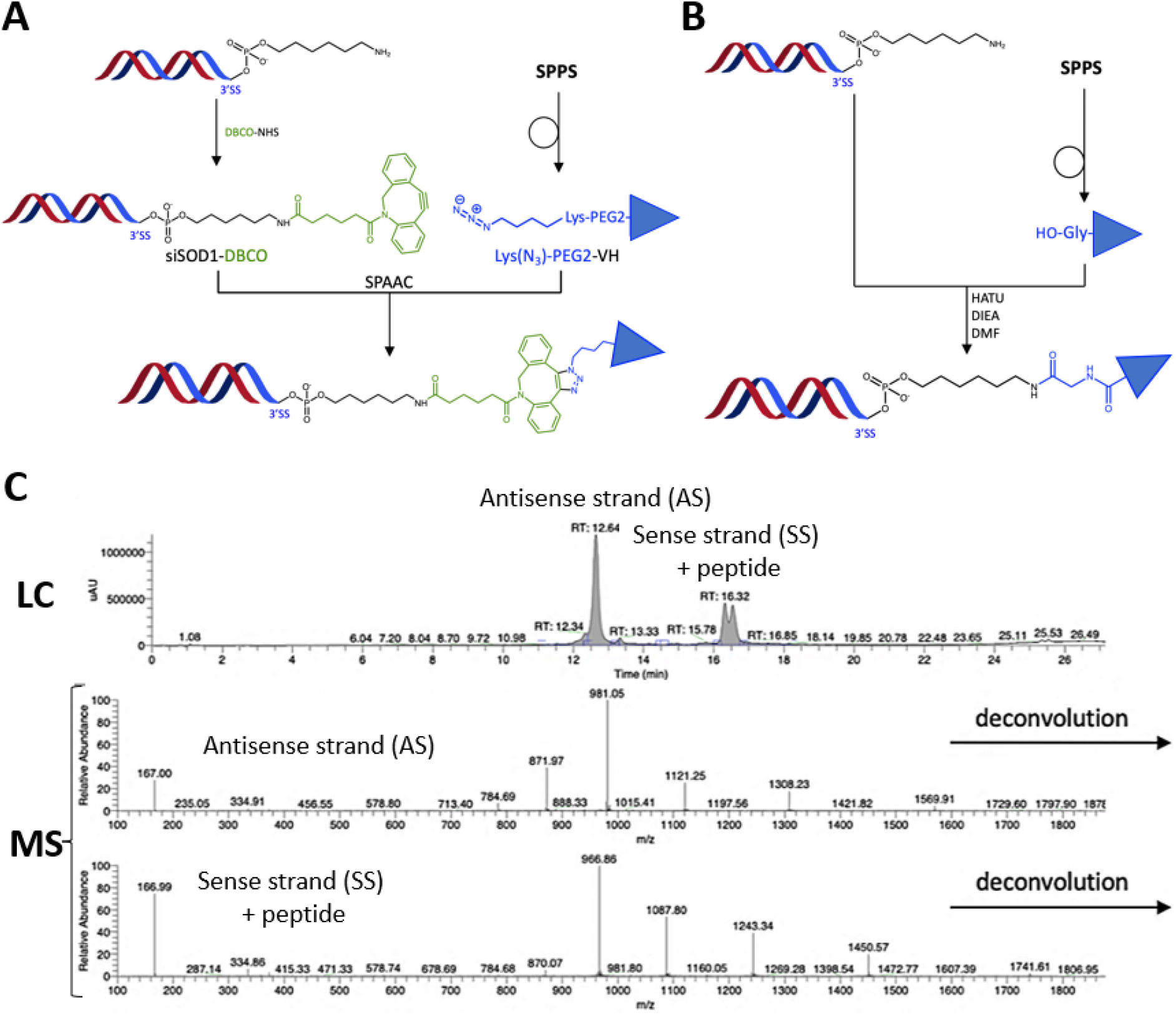
Synthesis strategies and characterization methods of siSOD1-peptide conjugates. **(A)** Production process of siSOD1-peptide conjugates. A constraint alkyne (DBCO) was introduced on the N6 modification at the 3’ end of siSOD1 sense strand. An azide function was incorporated in the VH4127 peptide sequence during SPPS in the form of azidolysine (Lys(N3) or K(N3)) and spaced from it with a PEG2. **(B)** Alternative synthesis strategy of siSOD1-peptide conjugates. The siSOD1-peptide conjugate was obtained through direct amidation between the N6 modification at the 3’ end of siSOD1m sense strand and the free carboxylic acid of peptide VH4127. **(C)** LC/MS characterization of siSOD1-peptide conjugates. Buffer A: HFIP 12.5mM and DIEA 4mM in H2O; Buffer B : HFIP 12.5mM and DIEA 4mM in MeOH; Flow rate was 0,3 mL/min and column temperature set at 65°C; Detection was done at 260 and 214 nm. MS analysis in negative mode and spectra was deconvoluted.

Among the different bioconjugation strategies, the SPAAC or copper-free “Click-Chemistry” reaction is particularly suitable for post-synthetic and site-specific conjugation of large biomolecules such as siRNAs^30^. Indeed, this reaction presents several advantages: i) the reactive functions involved are inert towards other chemical functions, conferring major reaction selectivity; ii) the reaction takes place at room temperature in both aqueous and organic solvents, thus facilitating the solubilization of reagents; iii) in opposition with the first version of this reaction, there is no need for copper as a catalyst, which is cytotoxic and particularly difficult to extract from the final product.

To prepare the siSOD1-peptide conjugates based on this click strategy, we chose to introduce the azide moiety onto the peptide and the alkyne moiety onto the siRNA. In this aim, the unnatural azidolysine bearing an azide function was additionally incorporated into the amino-acid sequence during the solid-phase peptide synthesis (SPPS). To retain the affinity of the VH4127 peptide towards the LDLR, the azidolysine was spaced out from the bulky cyclic peptide by introduction of a PEG2 linker. As for the constrained alkyne function, it was introduced post-synthetically by reaction of a heterobifunctional linker DBCO-NHS with the amine group of the hexylamino (N6) modification. The final click conjugation step allowed covalent attachment of the azido-peptide to the siSOD1-DBCO to obtain the siSOD1-peptide conjugate by SPAAC (Figure 2-A).

Additionally, to investigate the impact of the conjugate design on its biological properties, a direct conjugation method was investigated: the coupling without prior addition of the DBCO-NHS linker was achieved by a straightforward amidation between the siRNA’s N6 modification and the VH4127 peptide through its carboxylic acid C-terminal (Figure 2-B).

All crudes were purified by high-performance liquid chromatography (HPLC) with a >65% purity and characterized with HPLC-mass spectrometry (HPLC-MS). Molecular masses of both antisense and sense strands were calculated from a manual deconvolution. The HPLC conditions allowed separation of the two strands on column and consequently separate ionization in MS. All HPLC-MS analysis of siSOD1-peptide conjugates exhibited for the sense strand conjugated to the peptide moiety a double peak (UV) with the exact same mass. This double peak results from the formation of two regioisomers during the click-chemistry reaction step between the DBCO group and the azidolysine^31^. (Figure 2-C).

### LDLR-binding affinity of siSOD1-peptide conjugates using Surface Plasmon Resonance (SPR)

Chemical design of the conjugates may directly impact the affinity of the conjugates and thus modulate the productive uptake, intracellular trafficking and therefore biological or therapeutic effect^15,32^ of siRNA-peptide conjugates. We thus prepared a panel of different conjugates to investigate different parameters in the siSOD1 conjugate chemistry (Table 1). In general, the peptide conjugation at the 3’-end of the siSOD1 sense strand impacts only moderately its LDLR-binding affinity, with K_D_ values ranging from *c.a*. 10 to 100 nM, compared to the VH4127 peptide ligand alone with a *K_D_* of 40 nM. While the siSOD1 −33, −35, −36, −37 conjugates demonstrated the highest LDLR-binding affinity, the siSOD1-31 and −32 conjugates showed a slightly lower *K_D_*. More particularly, we observed that the presence of the 5’VP on the siSOD1-36 conjugate seems to positively influence receptor/ligand interaction in comparison with the siSOD1-32 conjugate, its homologue without the 5’VP modification. Interestingly, this finding was not observed with the siSOD1-37 and siSOD1-33 conjugates, respectively homologues with and without 5’VP, which exhibit LDLR-binding affinities in the same range. Peptide conjugation on the 5-LC-NU modification in these two conjugates, originally used to prevent interactions between a double-strand oligonucleotide and any large cargo linked to it, may explain this similarity by favoring ligand presentation on its target. Unfortunately, the siSOD1-34 conjugate could not be explored by SPR due to the strong complexation of the histidines contained in its structure with the nickel present on the SPR chips.

**Table 1:**
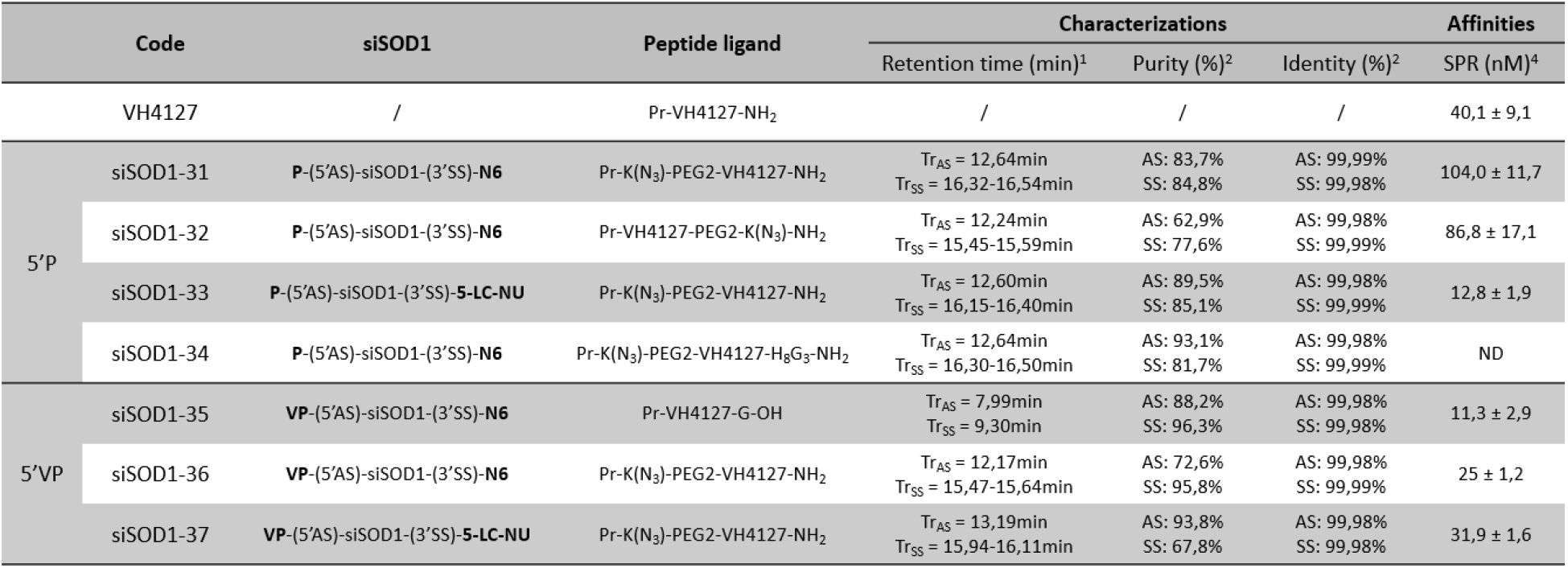
Physicochemical properties of siSOD1-peptide conjugates. **(1)**: Retention time obtained with the following RP-HPLC conditions: buffer A: HFIP 12.5mM and DIEA 4mM in H_2_O and buffer B: HFIP 12.5mM and DIEA 4mM in MeOH. Flow rate was 0,3 mL/min with a gradient of 5–40% B in 18 min. The only exception was for the siSOD1-35 conjugate that was characterized in 12 min. **(2)**: RP-HPLC-UV purity of both antisense (AS) and sense (SS) strands. **(3)**: Identity is calculated as follow: theoretical mass / experimental mass*100 **(4)**: Chip NiHC1000m (with imidazole); LDLR: 2100-2300RU; MCK mode; means ± SD n=2-5 independent experiment

**Table 2:**
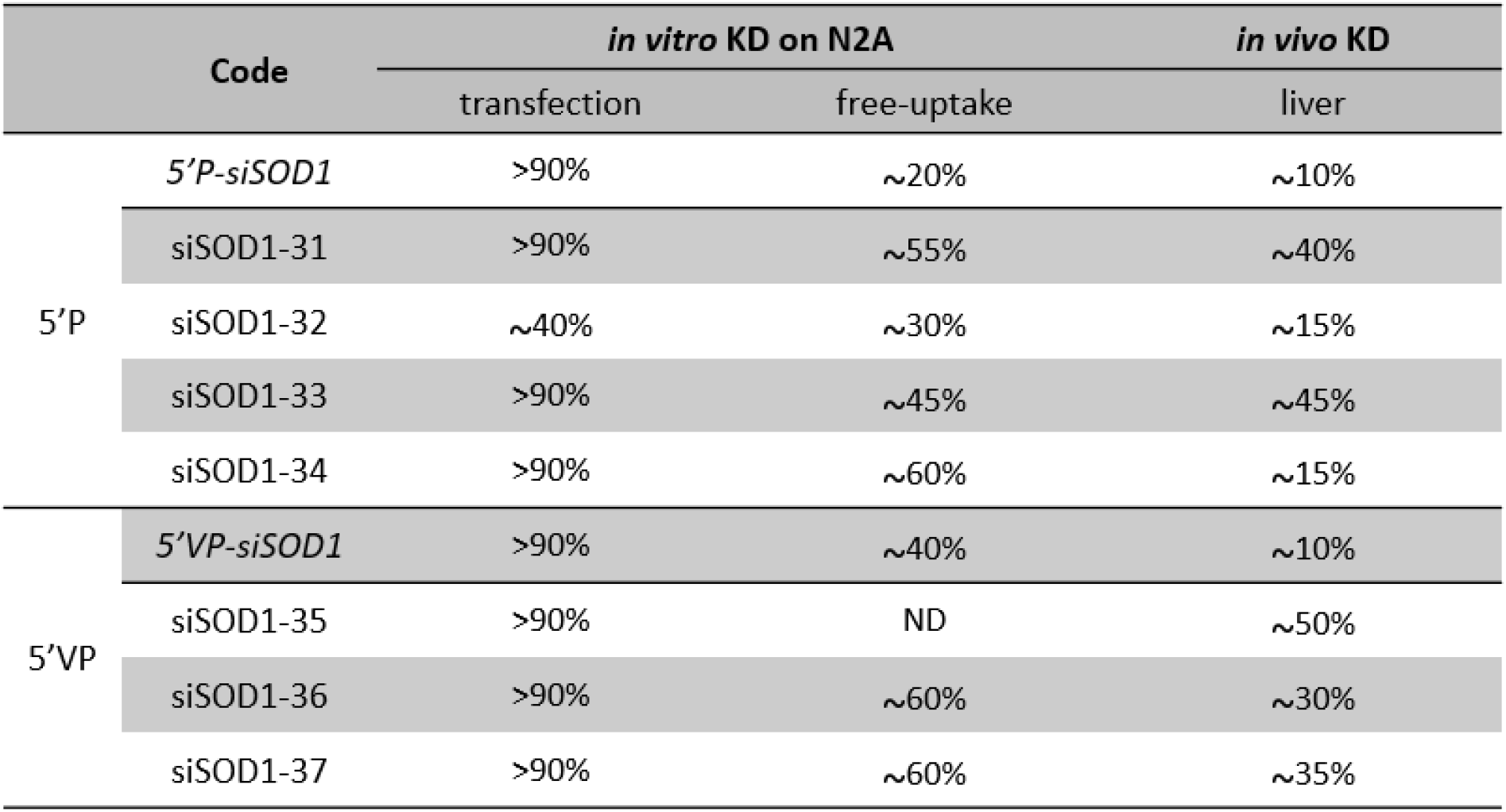
Summary of siSOD1-peptide conjugates *in vitro* and *in vivo* KD.

After verifying the good conservation of our peptide affinity post-conjugation, which is a prerequisite and the first key step for our conjugates to induce the RNAi response, we explored their gene-silencing efficacy *in vitro* on a Neuro2-A cell line.

### Gene-silencing potency of siSOD1-peptide conjugates in vitro in murine Neuro-2A cells

The *in vitro* gene silencing potential of siSOD1-peptide conjugates was investigated in the murine Neuro-2A (N2A) cell line after lipofection, to verify their intrinsic RNAi activity, or free uptake, to evaluate their potential to undergo LDLR-mediated functional uptake, leading to *mSOD1* target mRNA knock-down (KD). The ability of both LDLR ligands, including LDL particles (DiI-LDL) and the previously described LDLR-binding VH4127-A680 conjugate (vs. its non-binding scrambled version VH4sc-A680)^33^, to specifically bind the murine LDLR (mLDLR) expressed by N2A cells was verified beforehand (Supplemental Figure S1). First, transfection experiments clearly demonstrated that most of the tested siSOD1-peptide conjugates induced similar KD effect (*c.a*. 90%) to that of the unconjugated siSOD1 (Figure 4-A). The only exception was the siSOD1-32 conjugate (SPAAC with siRNA-N6 conjugation on the C-terminus of the peptide) with a knockdown potential of 40% following transfection. Second, free uptake experiments consistently demonstrated higher KD effect, up to ~60%, for most of the conjugates tested, compared to unconjugated siSOD1 molecules, demonstrating a higher functional uptake potential for LDLR-binding conjugates (Figure 4-B). Interestingly, whereas the unconjugated 5’VP-siSOD1 showed a slightly higher KD potential in this cellular model than the 5’P-siSOD1, with a mean of 40% vs. 22% respectively, this did not translate into a higher KD potential of 5’VP-siSOD1-peptide conjugates in the *in vitro* system. Again, the only exception among siSOD1-peptide conjugates was the siSOD1-32 conjugate, which did not show improvement compared to the unconjugated 5’P-siSOD1, consistent with its lower KD potential using lipofection. The results obtained with the 5’P-siSOD1-peptide conjugates also showed that the addition of a H_8_G_3_ poly-His stretch in C-ter of the VH4127 peptide, introduced to potentially increase endosomal escape^34^, did not translate into an improved KD effect, as compared to the structurally similar siSOD1-31 conjugate. Although we cannot rule out a higher dissociation from LDLR in early/sorting endosomes and thereby a higher delivery to late compartments, this may not result in higher endosomal escape and delivery to the cytosol. Finally, the minor KD effect of all tested conjugates comprising the non-binding scrambled peptide confirmed the involvement of LDLR in the functional uptake and KD effect of our siSOD1-peptide conjugates (Figure 4-C). We also investigated an additional conjugate where the VH4127 peptide was directly conjugated to the amino group of the 5’P-siSOD1-N6 precursor, leading to a conjugate with a much smaller linker. As observed with other conjugates, the resulting siSOD1-35 conjugate showed an improved KD effect compared to its non-binding control siSOD1-35sc. Altogether, these results prompted us to further investigate *in vivo* the KD potential of some of the LDLR-binding conjugates.

**Figure 4:**
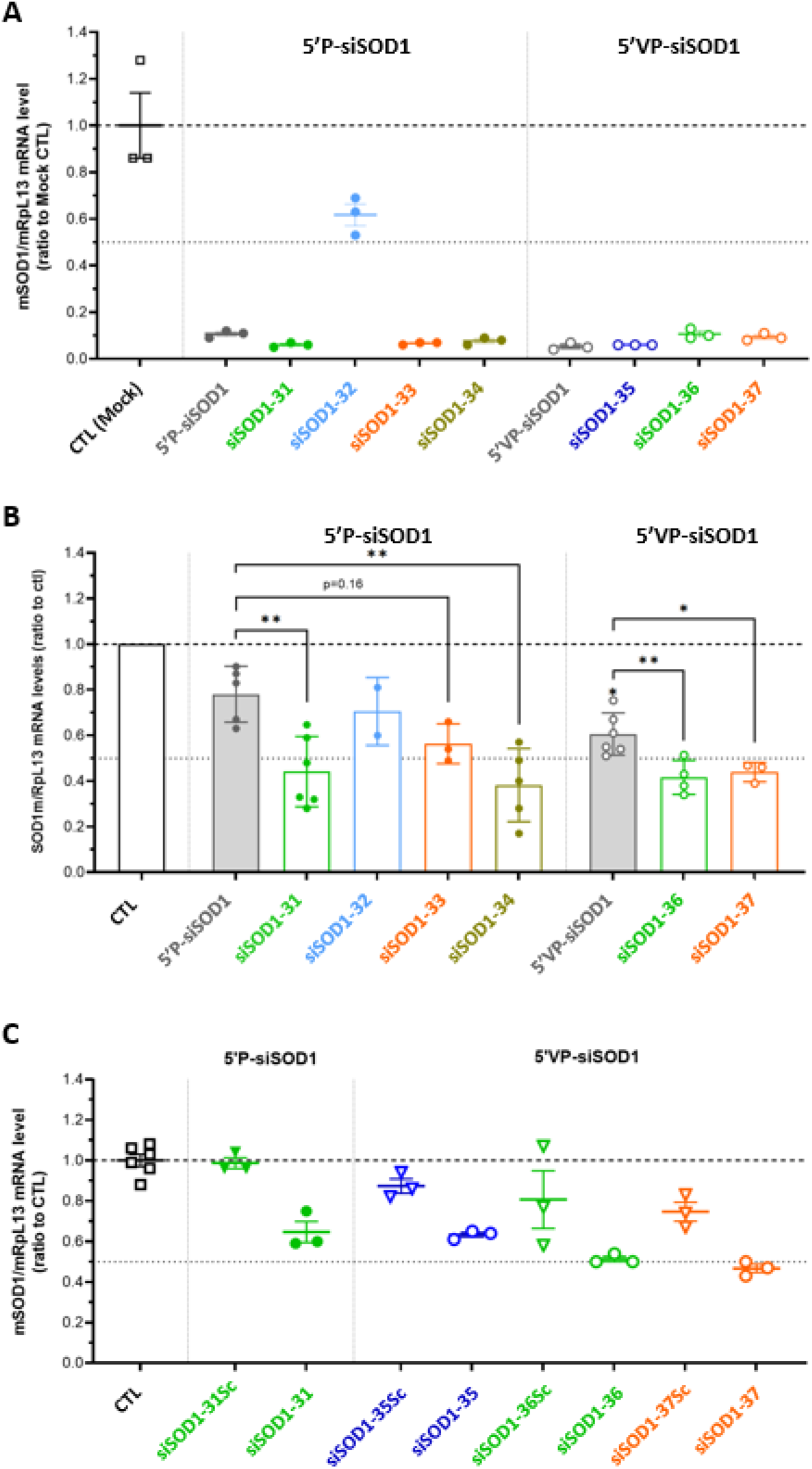
Gene-silencing potency of siSOD1-31, -32, -33, -34, -35, -36, -37 conjugates by transfection and free-uptake. **(A)** Transfection of siSOD1-peptide conjugates on Neuro-2a cells at 30nM. After 24h of incubation *SOD1* mRNA levels were amplified by RT-qPCR and quantified. **(B)** Free-uptake experiments of siSOD1-peptide conjugates on Neuro-2a cells at 1mM. After 3 days at 37°C, *SOD1* mRNA levels were amplified by RT-qPCR and quantified. Each dot corresponds to the mean value obtained in independent experiments. **(C)** Free-uptake experiments of siSOD1-31, -35, -36, -37 conjugates with their respective negative controls siSOD1-31Sc, -35Sc, -36Sc, -37Sc on Neuro-2a cells at 1mM. After 3 days at 37°C, *SOD1* mRNA levels were amplified by RT-qPCR and quantified.

### Gene-silencing potency of siSOD1-peptide conjugates after systemic administration in mice

The *in vivo* targeting and functional uptake potential of LDLR-binding siSOD1-peptide conjugates was evaluated as follows. Seven days after single intravenous (i.v. bolus) administration in mice at 1 μmole/kg (corresponding to 15 mg/kg siRNA), *SOD1* mRNA expression level was quantified using RT-qPCR in the liver, an organ expressing high levels of the target LDL receptor and where the VH4127 peptide previously demonstrated efficient and specific distribution^26,27^. As expected, both the unconjugated siSOD1 showed poor KD effect, even in the presence of the stabilizing 5’VP-AS modification (~10% KD, ns) (Figure 5). On the contrary, siSOD1 conjugation to the LDLR-targeting VH4127 peptide consistently led to a significant KD effect, reaching *c.a*. 50% KD with both siSOD1-33 and −35 conjugates. No clear correlation could be evidenced between *in vitro* free uptake and *in vivo* KD results. First, introduction of the 5’VP-AS modification in the unconjugated siSOD1, which was shown to increase both AS metabolic stability and RISC-engagement^35,36^, did not improve the liver KD effect as observed during our *in vitro* free uptake experiments (see Figure 4-B). Second, besides the siSOD1-32 conjugate that showed only minor, yet significant, KD effect (~20%, p<0.05) as observed in free uptake experiments, the siSOD1-33 conjugate with a 5’P-AS and the siSOD1-35 conjugate with a 5’VP-AS both demonstrated the highest KD potential, with a *c.a*. 50% reduction in *SOD1* mRNA levels (p<0.001), whereas the siSOD1-31, −34, −36 and −37 conjugates performed best in our *in vitro* setting.

**Figure 5:**
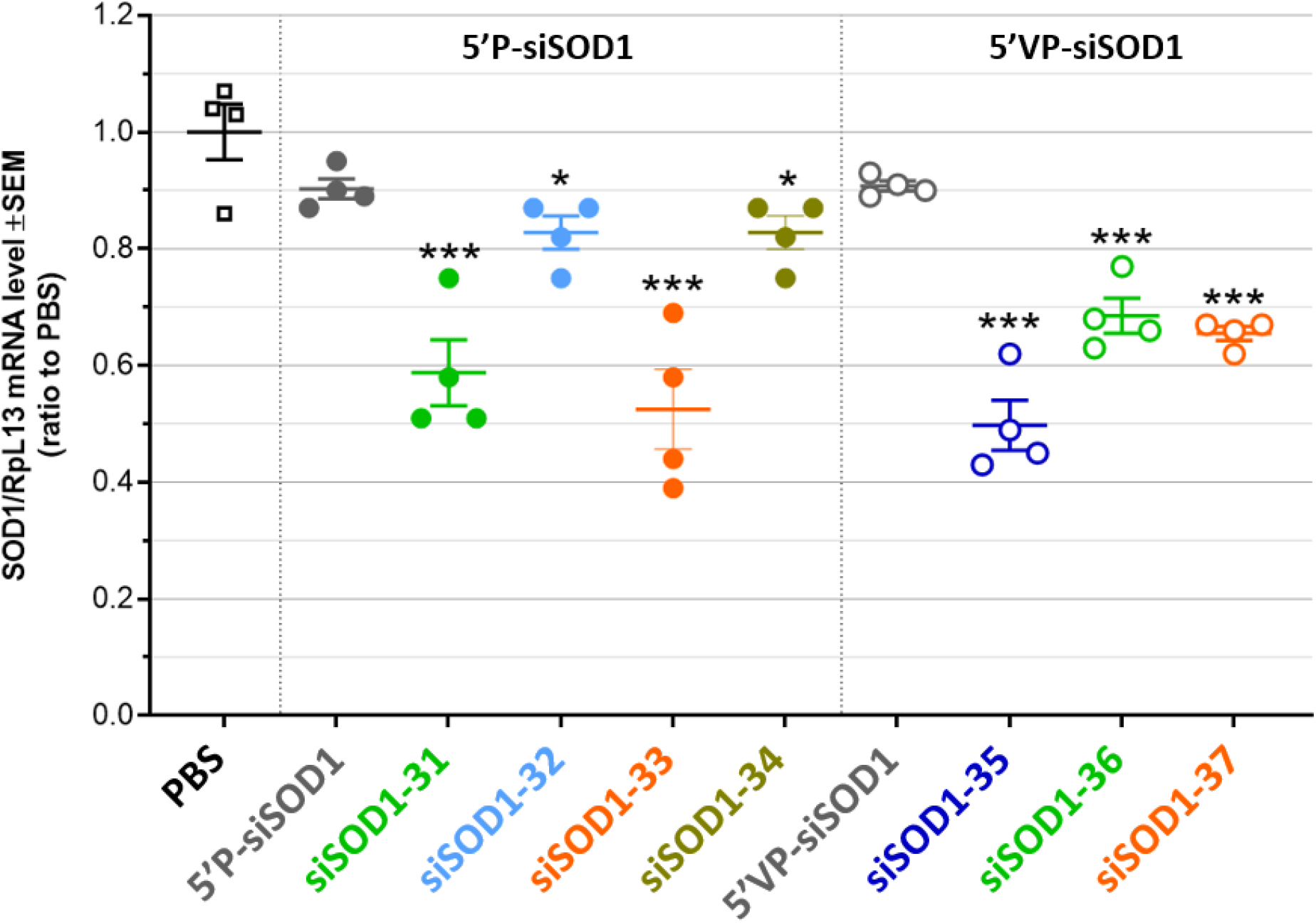
In vivo gene-silencing potency of siSOD1-31, -32, -33, -34, -35, -36, -37 in mice liver. Mice were injected (i.v. lateral tail vein) with a solution of siSOD1-peptide conjugate in PBS (15mg/kg). 7 days post-administration mice were euthanized and perfused with saline solution (0,9% NaCl). Organs were collected for *SOD1* RNAm amplification by RT-qPCR and quantification. Each dot corresponds to a mouse.

## DISCUSSION AND PERSPECTIVES

Many attempts have been made this last decade to overcome the inherent barriers to the clinical use of therapeutic siRNAs and establish the necessary foundations for the development of strategies that allow their functional delivery to a tissue of interest. With more than 30 years of efforts on the chemistry and stabilization of siRNAs it is now possible to produce molecules containing chemical modifications to achieve high metabolic stability, high target sequence specificity, and efficacy. Nevertheless, exploiting the full clinical potential of siRNAs, which could potentially make the term “undruggable” obsolete ^37^, still requires the development of efficient systems to address major delivery hurdles including fast plasma clearance, low tissue selectivity and low functional uptake in target cells. In the present work, we validated the potential of a previously described LDLR-binding cyclic peptide ^26,27,33^ for the *in vitro* and *in vivo* targeting and productive delivery of therapeutic siRNAs. Although no clear *in vitro*-to-*in vivo* structure-activity relationship could arise for our LDLR-targeting siSOD1-peptide conjugates, as discussed thereafter, it clearly appeared that their LDLR-binding potential translates into an active functional uptake in cells in vitro and *in vivo* in the liver, leading to significant RISC engagement and KD effect.

As shown in an early study by Gilleron et al., where small molecules and different delivery systems based on either LNPs or molecular cholesterol-siGFP conjugates were screened for their functional uptake potential, improvement of the gene silencing occurred on either the uptake system *per* se, or on subsequent trafficking and endosomal escape steps ^38^. RNAi induced by siRNA-ligand conjugates occurs in four basic steps: i) the effective conjugate recognition by the targeted receptor, without disturbing the endogenous ligand interaction; ii) the endocytosis of the siRNA-ligand/receptor complex within cells, followed by conjugate dissociation from receptors into early and/or sorting endosome; iii) the siRNA endosomal escape to reach the cytoplasmic compartment; iv) its recognition and loading into the RISC complex.

In the present study, we explored the potential of LDLR-binding peptides to efficiently transport siRNA through these prerequisite steps, and to evaluate whether the modulation of some of the siRNA-peptide conjugate parameters could impact their efficacy. The LDLR represents an attractive cell-surface target receptor to support efficient functional delivery of therapeutic oligonucleotides into different cells and organs, owing to its relatively high cell surface expression, with some degree of tissue selectivity, and its ability to undergo hundreds of endocytosis cycles during its 20-hour lifespan^22,27,39,40^. However, it is worth noting that efficient functional delivery of an ASO to beta pancreatic cells in mice could be achieved by means of a GLP-1 synthetic peptide ligand targeting the GLP1R, a cell-surface receptor from the GPCRs superfamily which are known for their rather low expression and low endocytic capacity^41,42^. Therefore, beyond the initial assumption that delivering therapeutic oligonucleotides to extrahepatic sites requires new Ligand/Receptor pairs displaying features similar to those of the prototypical GalNAc/ASGPR pair, it appears that other parameters can largely compensate for poor expression and endocytic potential. In line with this, the molecular mechanisms underlying and enabling the mostly inefficient (below 1%) transit of Ligand-Oligo conjugates from early endocytic vesicles to the cytosol or nucleus where the oligo can eventually engage its target, remain poorly understood^43–46^. For these reasons and given the unique features of each new ligand/receptor pair, investigation of the structure-activity relationship underlying Ligand-Oligo activity, including in the present work, is performed for now with a rather empirical approach^47^.

A siSOD1 was studied based on the stabilization scheme described by Foster et al. It encompasses chemical modifications that increased its resistance to nucleases and silencing potency while diminishing potential off-target effects and cytotoxicity^29^. We selected the N6 and the 5-LC-NU modifications as reactive functional amino groups on the 3’ end of the sense strand for further conjugation with our peptide ligands. Antisense strand 5’end modifications are also known to increase siRNA stability and potentiate their effects presumably via enhanced loading within the RISC complex. Thus, we also compared conjugates encompassing a 5’-phosphate or 5’-vinylphosphonate modification on the siRNA antisense strand.

From a general medicinal chemistry perspective, conjugation modality and linker chemistry used so far in recent Ligand-oligonucleotide conjugates involve generally either direct amidation coupling or SPACC.

Examples of these conjugation strategies include siRNA-GalNac conjugates^48^, ASO-peptide conjugates^42,47,49^, siRNA-antibody conjugates^50^. SPAAC offers high coupling yields and versatility, allowing linkage of complex biomolecules in a highly specific manner. Nonetheless, the bulky and hydrophobic nature of the constrained alkyne precursor (DBCO, BCN) can possibly influence the conjugate tridimensional structure, and thus negatively impact its physico-chemical proprieties. On the contrary, direct amidation coupling allows a more straightforward coupling and generates a much smaller, natural, hydrophile, and flexible linker. However, this coupling strategy is less versatile and requires two highly reactive chemical partners, namely a primary amine and a carboxylic acid, thus obliging careful upstream considerations on the synthesis strategy in which it is applied.

Considering the bulky and charged nature of our peptide ligand, consisting of a constrained cyclic octapeptide comprising two arginine residues, and its use as a targeting ligand for a covalently attached siRNA, the modulation of its anchoring site as well as the linker length and nature can have profound impact on its orientation and accessibility for optimal binding to the LDLR. In the present work, different siSOD1-peptide conjugates were generated with the objective to investigate how these parameters can impact both LDLR-binding and siSOD1-peptide knock-down potential in both a cellular model and after systemic administration in mice. Interestingly, with respect to their LDLR-binding potential, our results show rather low impact of conjugation site and chemistry on *K*_D_ values compared to the unconjugated peptide (40nM), demonstrating a permissive peptide/LDLR interaction even in the presence of the conjugated siRNA. Indeed, variation of parameters such as i) the peptide coupling site (N-ter vs. C-ter), ii) the sense strand 3’-end coupling site with the 5-LC-NU modification (Figure 1-B) or iii) the linker length and nature, much smaller and less hydrophobic produced by direct coupling (Figure 2-B), did not induce drastic increase or loss of LDLR-binding affinity which remained in the range of 10 to 100 nM. Therefore, one can anticipate broad application potential of our LDLR-targeting ligand in terms of oligonucleotide modality, such as gapmer ASOs, ssASOs, etc. On the contrary, other ligand/receptor pairs investigated for oligonucleotides targeted delivery purposes demonstrated altered binding and uptake by their target receptor, attributed to some interference between the negatively charged oligonucleotide backbone and positive charges of the ligand. These include the neurotensin (NT) peptide proposed as a targeting ligand for improved delivery to NTR-expressing neurons^51^, or the Glucagon-like Peptide 1 (GLP1) that targets the GLP1R expressed by pancreatic beta cells^47,52^.

After demonstrating adequate binding of our siSOD1-peptide conjugates to cell-surface LDLR, the different coupling strategies proposed in the present work could impact downstream events. One of the most critical steps following receptor-mediated endocytosis relies in peptide/LDLR dissociation in sorting endosomes and subsequent creation of a late endosomal or lysosomal depot, a prerequisite for siRNAs to slowly egress in the cytosol where they can engage the RISC. Although we could not verify the efficient uptake and intracellular accumulation of the siSOD1-VH4127 conjugates of the present study, we demonstrated using a pulse-chase procedure with a fluorescent A680-VH4127(3’SS)-siGFP conjugate the efficient delivery from LDLR-positive early compartments to LDLR-negative late compartments (data not shown). As performed by most studies in the field, these intracellular trafficking steps were investigated indirectly, by measuring the final functional read-out of our SOD1-peptide conjugates, namely the *SOD1* mRNA knock-down, in both a cellular setting and after systemic administration in mice. Importantly, our results clearly demonstrate a significant improvement of siSOD1 functional uptake and gene knock-down, when conjugated to the LDLR-targeting VH4127 peptide, both in murine Neuro-2a cells expressing functional LDLR and *in vivo* in the liver of mice injected with our conjugates, with *c.a*. 50% reduction in *SOD1* mRNA levels with the best conjugates, while only minor effect could be observed with the unconjugated siSOD1. These results thus demonstrate that the LDLR-binding peptide not only maintains its binding potential when conjugated to a siRNA, but also mediates efficient uptake leading to significant gene knock-down both *in vitro* and *in vivo*. However, we observed a discrepancy in the relationship between the conjugation strategy and the knock-down efficiency in either our cellular model or in mice. The best performing conjugates after systemic administration in mice were the siSOD1-33 and the siSOD1-35 conjugates. In the siSOD1-33 conjugate, the VH4127 peptide is coupled *via* an N-terminal Lys(azido)-PEG2-group to a pseudo-nucleotide (5-LC-NU) extending from the 3’-end of the sense strand of a 5’P(AS)-siSOD1. The siSOD1-35 conjugate was obtained by direct coupling of the VH4127 *via* a C-terminal -Gly-OH group to an hexylamino group protruding from the 3’-end of the sense strand of a 5’VP(AS)-siSOD1. On the contrary, these two conjugates showed rather low functional uptake potential upon free uptake in Neuro-2a cells compared to other LDLR-binding conjugates. First, the fast clearance rate of our siSOD1-peptide conjugates that may occur after systemic (i.v. bolus) administration in mice may lead to much shorter exposure duration of target cells, compared to continuous 3-day exposure of N2A cells in our cellular model. This in turn could favor the siSOD1-33 and −35 conjugate from a plasma pharmacokinetics perspective in a way that remains unclear. Second, liver hepatocytes *in vivo* vs. cultured murine neuroblastoma cells can display profound differences in their intracellular dynamics, making some molecular designs more prone to reaching the cytosolic RISC complex in pharmacological amounts.

Recent studies showed that siRNA accumulation and stability in acidic intracellular compartments is critical for long-term activity^53^, and that fewer than 1% of endocytosed molecules reach the cytosol compartment^21,41,43,45,54^. Therefore, beyond the identification of new ligand/receptor pairs able to support efficient functional uptake of therapeutic oligonucleotides, endosomal escape remains a rate-limiting step for oligonucleotide functional delivery and there is a high need to investigate new strategies to increase the amount of molecules reaching the cytosol^55^. Although several methods have been investigated, based on either osmolytic compounds such as chloroquine or membrane-destabilizing agents such as peptides derived from the influenza hemagglutinin (HA), they all suffer from toxicity concerns making them non-viable approaches for *in vivo* applications^46,53,55,56^. In the present work, we investigated an original approach by introducing a pH-conditional endosomal escape-inducing peptide (EEIP) directly conjugated in C-terminal of our VH4127 peptide. Because the lateral imidazole group of histidines have a pK of 6.0, corresponding to the pH found in early and sorting endosomes, they can remain neutral at extracellular pH while protonated when reaching these endosomal compartments following receptor-mediated endocytosis. Several studies have shown that poly-histidine stretches can act as EEIPs by gaining the ability to translocate across its monolayer membrane, thereby improving the KD potential of the conjugated siRNA^34,57^. We have confirmed that conjugation of a 8-histidine stretch in C-terminal of our LDLR-binding VH4127 peptide led to a strong increase in intracellular delivery in different *in vitro* cellular models, and that this translates into a 2- to 3-fold increase in LDLR-enriched tissue exposure after systemic administration in mice (data not shown). In the present study, the siSOD1-34 conjugate encompassing the same linear 8-histidine stretch in C-terminal of the VH4127 peptide was produced to explore its potential to increase the functional delivery of the siRNA. Unfortunately, the LDLR-binding affinity of this conjugate could not be evaluated by SPR due to the strong complexation of histidine with the nickel present on the chips. Despite promising results in transfection, this conjugate did not show higher knock-down effect *in vitro* and even showed a lower knock-down effect *in vivo*. As suggested from positively-charged peptide ligands or cell-penetrating peptides (CPPs)^51,55^, unwanted molecular interactions occurring in sorting and late endosomes between the positively charged poly-histidine moiety and the negatively charged siRNA backbone could mask the protonated poly-His stretch and thus hamper the expected benefit in late endosomal delivery and/or endosomal escape potential, which in turn might explain the observed drop of efficacy ^54^.

Finally, even if the conjugates enter cells and undergo endosomal escape their ability to load into the RISC depends on the presence of a phosphate group at the 5’-end of the active/antisense strand^58^. Because this functionally crucial spot might be cleaved off by lysosomal phosphatases, introduction of the 5’VP modification represents a suitable alternative to 5’P, with a potential to improve our conjugate silencing effect^35,36^. The siSOD1 −35, −36, and −37 conjugates were produced with a 5’VP instead of a 5’P to investigate this aspect. None of these conjugates showed a higher knock-down effect than with the 5’P-containing conjugates, both in free uptake experiments and *in vivo*. One possible explanation is that the expected benefit from this 5’VP modification, namely higher metabolic resistance in lysosomal compartments, cannot occur due to insufficient delivery to these compartments. In this hypothesis, the rate-limiting step of the LDLR-binding siSOD1-peptide conjugates might rely on an earlier step during intracellular trafficking, such as low dissociation from LDLR in sorting endosomes leading to recycling back to the cell surface (non-productive uptake).

The present work demonstrates that our LDLR-targeting peptides can support efficient functional uptake of a model siRNA in both *in vitro* and *in vivo* settings. However, we need to gain further understanding of the rate-limiting steps and identify strategies to further improve the conjugate design to mediate significant knock-down effects at lower concentrations and doses. One possible approach includes the use of a fluorescent probe, such as the A680, conjugated at the 3’-end of the active/antisense strand, allowing direct tracking in cellular models and evaluation of the intracellular fate of tested conjugates. We previously used approach to visualize early recycling vs. lysosomal trafficking pathways and quantify the cellular elimination profile of fluorescent conjugates^33^. Another strategy relies on the use of a GalNAc-INF7 conjugate as performed previously^53^ to force endosomal escape of potential intravesicular depot of an oligonucleotide in liver hepatocytes, if any. In this approach, any improvement of the knock-down effect following GalNAc-INF7 injection would confirm that the oligonucleotide was efficiently delivered in cells but with endosomal escape being the rate-limiting step towards optimal RISC-engagement and knock-down effect. On the contrary, no improvement would indicate insufficient intravesicular depot and guide further design optimization towards higher uptake and dissociation from the LDLR in early/sorting endosomes.

## Conclusion

Here, we validate the previously described synthetic LDLR-binding peptides, including the VH4127 peptide, as viable ligands able to trigger efficient LDLR-mediated functional delivery of therapeutic oligonucleotides both in a cellular model of neuroblastoma and *in vivo* after systemic administration in mice. Although no clear *in vitro*-to-*in vivo* functional correlation could be made from these two distinct settings, our results highlight several potential tracks for further optimization of this new class of Ligand-oligo conjugates. Since the LDLR presents some degree of tissue selectivity, beyond the liver, and is overexpressed in many cancers^22,27,39,40^, the present work opens new opportunities and warrants further evaluation for delivery to either extrahepatic sites or in tumors that otherwise do not support functional uptake of naked oligonucleotides.

## MATERIAL AND METHODS

### Chemical reagents and Material

Fmoc-amino acids were supplied from Iris Biotech (Marktredwitz, Germany). All other amino acids and reagents were purchased from Sigma-Aldrich or Analytical Lab. Peptide assembly was carried out using the Liberty (CEM^®^) microwave synthesizer by solid phase peptide synthesis (SPPS) in Fmoc/tBu strategy. Fmoc-Rink amide aminomethylpolystyrene resin (loading 0.74 mmol/g) and Fmoc-Gly-Wang resin (loading 0.66 mmol/g) were purchased from Iris Biotech was used as solid support. All siRNA were purchased from Horizon Discovery or Genelink.

### Purification and Analytical Methods

#### Peptides-based products

Monitoring of reactions and quality controls of the peptide-based intermediates were carried out by LC/MS. The LC/MS system used was a Thermo Scientific Ultimate 3000 liquid phase system equipped with an ion trap (LCQ Fleet) and an electrospray ionization source (positive ion mode). The LC flow was set to 2mL/min using H_2_O_0.1%TFA_ (buffer A) and MeCN_0.1%TFA_ (buffer B) as eluents. The gradient elution was 10-90% B in 5min (monitoring) or 10min (quality control). The heated electrospray ionization source had a capillary temperature of 350°C. Crude peptides were purified by RP-HPLC on a Thermofisher UltiMate®3000 system equipped with a C18 Luna™ (5 μm, 100 mm × 21.2 mm). Detection was done at 214 nm. The elution system was composed of H_2_O_0.1%TFA_ (buffer A) and MeCN_0.1%TFA_ (buffer B). Flow rate was 20 mL/min.

#### Oligos-based products

Monitoring of reactions and quality controls of the intermediate and final products were carried out by LC/MS. The LC/MS system used was a Thermo Scientific Ultimate 3000 liquid phase system equipped with an ion trap (LCQ Fleet) and an electrospray ionization source (negative ion mode). The LC flow was set to 0,3mL/min using HFIP 12.5mM and DIEA 4mM in H_2_O (buffer A) and HFIP 12.5mM and DIEA 4mM in MeOH (buffer B) as eluents. The gradient elution was 5–40% B in 18 min and the column temperature set at 65°C. The heated electrospray ionization source had a capillary temperature of 350°C.

### Sequence and Modifications of siRNAs

siSOD1 sequences and modifications for this study were as follow: siSOD1 sense strand = 5’-P.mC.*.mA.*.mU.mU.mU.mU.2’-F-A.mA.2’-F-U.2’-F-C.2’-F-

C.mU.mC.mA.mC.mU.mC.mU.mA.mA.mA.N6-3’, antisense strand = 5’-P.mU.*.2’-F-

U.*.mU.mA.mG.2’-F-A.mG.2’-F-U.2’-F-G.mA.mG.mG.mA.2’-F-U.mU.2’-F-

A.mA.mA.mA.mU.mG.*.mA.*.mG-3’; where mN and 2’-F-N represent 2’-O-methyl and 2’-Fluoro sugar-modified RNA nucleosides, respectively. P represents 5’-phosphate ending, VP represents 5’-vinylphopshonate, N6 represents the 6-carbon aliphatic arm finishing by an amine and 5-LC-NU represents the modified nucleoside 5-aminohexylacrylamino-uridine bearing a 6-carbon aliphatic arm at the position 5 of the uridine. Finally, * represents a phosphonothioate linker (PS). All the siRNAs were synthesized by Horizon Discovery and Genelink, desalted, and dried-out without further purification.2/3/2023 9:46:00 AM

### Preparation of peptide ligand precursors

Peptides Pr-K(N_3_)-PEG2-[<u>cMThzRLRGPen</u>]_c_-NH_2_; Pr-[<u>cMThzRLRGPen</u>]_c_-PEG2-K(N_3_)-NH_2_; Pr-[<u>cMThzRLRGPenG</u>]_c_-OH; Pr-K(N_3_)-PEG2-[<u>cMThzRLRGPen</u>]_c_-H_2_GH_2_GH_2_GH_2_-NH_2_ where “c” indicates a cyclic peptide, were synthesized by SPPS under microwave activation. Briefly, Fmoc-Rink amide aminomethylpolystyrene for C-terminal amide peptides or Fmoc-Gly-Wang resin for C-terminal acid peptides were swollen in DMF for 10min. Initial deprotection of the resin and stepwise assembly of the Fmoc-protected amino acids were performed under microwave activation using standard Fmoc/tBu peptide chemistry. Coupling times of 300 sec were used with solution of Fmoc-amino acid (1 eq), Oxyma (10 eq excess; 1M) in DMF, and DIC (5 eq excess; 1M) in DMF under micro-wave activation at 70°C. Fmoc removal was carried out with piperidine/DMF (20:80 v/v) for 200 sec under micro-wave activation at 75°C. Double coupling was used for Arg and Met amino acids to improve yielding. In the special case of (D)-cysteine, coupling was performed at 50°C for 360 sec to avoid unwanted racemisation. Finally, DIC (5 eq excess; 1M) in DMF under micro-wave activation at 70°C. Fmoc removal was carried out with piperidine/DMF (20:80 v/v) for 200 sec under micro-wave activation at 75°C. Double coupling was used for Arg and Met amino acids to improve yielding. In the special case of (D)-cysteine, coupling was performed at 50°C for 360 sec to avoid unwanted racemisation. Finally, the peptidyl-resin was washed successively with DCM, MeOH and DCM. N-terminus propionylation on solid support was carried out manually in a Pr_2_O/DCM (1:1) mixture for 5 min twice. The resin was further washed three times with DCM. and then treated with TFA/TIS/H_2_O (95:2,5:2,5) containing DTT (100mg/mL) at room temperature for 2 h. The cleavage solution was recovered, concentrated under N_2_ flow, and precipitated three times in cold diethyl ether. The crude products were dissolved in an H_2_O_0.1% TFA_ /CH_3_CN_0.1% TFA_ (1:1) mixture and lyophilized. To form the disulfide bridge between the first D-Cys residue and the mixture and lyophilized. To form the disulfide bridge between the first D-Cys residue and the last Pen residue, peptides were then dissolved in 0,5% aqueous AcOH (concentration of 0,5 mg of peptide per mL). pH of the peptide solution was adjusted to 8-9 with 2M (NH_4_)_2_CO_3_ and K_3_Fe(CN)_6_ was added as a mild oxidative agent at room temperature for 0,5 h. The crude products were directly purified by preparative RP-HPLC according to the method described previously. Final products were characterized to assess their purity and identity by HPLC-MS.

| Peptide sequences | RP-HPLC, retention time gradient | HPLC-UV purity (%) | Theoretical mass (Da) | ESI-MS (m/z) [M,H]<sup>+</sup> (calculated) |
| --- | --- | --- | --- | --- |
| Pr-K(N<sub>3</sub>)-PEG2-[cMThzRLRGPen]<sub>c</sub>-NH<sub>2</sub> | rt=3,36min | 91,7% | 1332,60Da | 1333,51Da |
| Pr-[cMThzRLRGPen]<sub>c</sub>-PEG2-K(N<sub>3</sub>)-NH<sub>2</sub> | rt=3,61min | 96,7% | 1332,60Da | 1333,00Da |
| Pr-[cMThzRLRGPenG]<sub>c</sub>-OH | rt=2,90min | 97,0% | 1091,45Da | 1092,69Da |
| Pr-K(N<sub>3</sub>)-PEG2-[cMThzRLRGPen]<sub>c</sub>-H<sub>2</sub>GH<sub>2</sub>GH<sub>2</sub>GH<sub>2</sub>-NH<sub>2</sub> | rt=2.68min | 90,5% | 2244,95Da | 2245,44Da |

### Preparation of siSOD1-peptide conjugates

#### siSOD1-peptide conjugate synthesis by SPAAC

The siSOD1 duplex (1eq) with a N6 or 5-LC-NU modification at the 3’-ending of the sense strand were dissolved in an H_2_O/DMF (1:1) mixture to reach a concentration of 0,5mM, in which DIEA (50eq) was added. Then, DBCO-NHS (10eq; 16,7mM) was dissolved in DMF and added to the siSOD1 solution. The reaction was allowed to stir at room temperature for 1,5h and followed-up by RPLC-MS. When the reaction was complete, the siSOD1-DBCO was precipitated in three times the reaction volume of cold absolute ethanol (EtOH) and resuspended in water. Then, the peptide ligand containing an azidolysine was dissolved in a small volume of DMF and added to the siSOD1-DBCO solution under magnetic agitation. The reaction was stirred for 1 h at room temperature and followed-up by HPLC-MS until completion. The crude products were purified by precipitation in cold absolute EtOH or by filtration on Amicon 3K, the final products were characterized by HPLC-MS and finally quantified by optical density at 260nm.

#### siSOD1-peptide conjugate synthesis by direct amidation

To a solution of peptide Pr-[<u>cMThZRLRGPenG</u>]_c_-OH in anhydrous DMF (20eq; 1,68mM) were successively added a solution of DIEA in anhydrous DMF (80eq; 80,5mM) and a solution of HATU in anhydrous DMF (20eq; 20,16mM) for pre-activation of the C-terminal acid. Pre-activation was allowed to proceed at RT for 15min and then at 40°C for 5min. Then, a solution of siSOD1-(3’SS)-N6 in H_2_O (1eq; 6mM) was added to the preactivated peptide and allowed to stir at 40°C for 1h. Monitoring of the reaction was performed by LC/MS. In the case of incomplete reaction after 1h, a second addition of the preactivated peptide was performed in the same conditions as described above. The crude products were purified by precipitation in cold absolute EtOH, the final products were characterized by HPLC-MS and finally quantified by optical density at 260nm.

|  | Sense strand + peptide |  |  | Antisense strand |  |  |
| --- | --- | --- | --- | --- | --- | --- |
|  | RP-HPLC, retention time gradient | Theoretical molecular weight | ESI-MS (m/z) [M,H] (calculated) | RP-HPLC, retention time gradient | Theoretical molecular weight | ESI-MS (m/z) [M,H] (calculated) |
| **siSOD1-31** | 16,32min<br>16,54min | 8709,66Da | 8710,80Da<br>8710,65Da | 12,64min | 7855,17Da | 7855,73Da |
| **siSOD1-31Sc** | 14,18min<br>14,33min | 8662,56Da | 8663,22Da<br>8663,03Da | 11,44min | 7855,17Da | 7854,96Da |
| **siSOD1-32** | 15,45min<br>15,59min | 8709,66 | 8709,80Da<br>8709,60Da | 12,24min | 7855,17Da | 7855,70Da |
| **siSOD1-33** | 16,15min<br>16,40min | 9004,91Da | 9005,42Da<br>9005,29Da | 12,60min | 7855,17Da | 7856,04Da |
| **siSOD1-34** | 16,30min | 9977,95Da | 9978,06Da | 12,64min | 7855,17Da | 7856,19Da |
|  | 16,50min |  | 9978,08Da |  |  |  |
| **siSOD1-35** | 9,30min | 8166,97Da | 8166,66Da | 7,99min | 7851,18Da | 7850,70Da |
| **siSOD1-35Sc** | 8,12min | 8088,89Da | 8088,74Da | 8,03min | 7851,18Da | 7850,79Da |
| **siSOD1-36** | 15,47min<br>15,64min | 8709,66Da | 8710,05Da<br>8710,25Da | 12,17min | 7851,18Da | 7851,35Da |
| **siSOD1-36Sc** | 13,56min<br>13,98min | 8694,86Da | 8694,86Da<br>8694,20Da | 11,47min | 7851,18Da | 7850,92Da |
| **siSOD1-37** | 15,94min<br>16,11min | 9004,91Da | 9004,75Da<br>9004,89Da | 12,19min | 7851,18Da | 7850,92Da |
| **siSOD1-37Sc** | 14,38min<br>14,49min | 8941,94Da | 8942,50Da<br>8942,35Da | 11,51min | 7851,18Da | 7850,86Da |

### Surface Plasmon Resonance

His-tagged recombinant human LDLR (extracellular domain, hLDLR-ECD) was purchased from SinoBiological (Beijing, China). SPR measurements were performed at 25 °C using a Biacore T200 apparatus (GE Healthcare) and 50 mM HEPES-NaOH pH 7.4, 150 mM NaCl, 50 μM EDTA, 0.005% Tween-20 (v/v), 10 mM Imidazole as running buffer. hLDLR was immobilized on a NiHC1000m sensor chip (Xantec, Dusseldorf, Germany) at a density of around 33 fmol/mm2. A control flowcell without hLDLR was used as reference. The multiple-cycle kinetic (MCK) method was applied to study molecular interactions of peptides and conjugates. Five different concentrations of the analyte were prepared by two-fold dilutions in running buffer and serially injected over the flowcells during 120 s at 30 μL/min with a dissociation time of 600 s after the last injection. Blank runs of running buffer were performed in the same condition. Double-subtracted sensorgrams were fitted globally with the 1:1 Langmuir model using Biacore T200 Evaluation version 2.0 software. Equilibrium dissociation constant (*K*_D_) are summarized in Table 1. All data are means of two to six independent experiments, each resulting from triplicate analysis.

### Cell culture and reagents

Neuro-2a cells were provided from ECACC and cultured in DMEM High Glucose GlutaMAX™ (DMEM) supplemented with 10 % fetal calf serum, 100μg/mL streptomycin, 100U/mL penicillin in a humidified 5 % CO2 atmosphere at 37 °C.

Dubelcco’s Phosphate Buffer Saline (D-PBS), Phosphate Buffer Saline no calcium no magnesium (PBS), DMEM, fetal calf serum, Penicilin/Streptomycin, 0.05% Trypsin/EDTA (Trypsin), EDTA, DiI-LDL were purchased from ThermoFisher.

#### In vitro validation of LDLR-expressing Neuro-2a cells

Neuro-2a cells were plated in a 96-well plate at 40 000 cells/well two days before the experiment. Cells were incubated 3 hours at 37°C with red fluorescent DiI-LDL particles at 20μg/mL, fluorescent cargo peptide A680-VH4127 or cargo scramble peptide A680-VH4Sc both at 1μM; or co-incubated with DiI-LDL particles and A680-VH4127 or A680-VH4Sc at the same concentrations than previously described. Cells were extensively washed in D-PBS then scrapped using Trypsin during 5min at 37°C and pelleted by centrifugation for 5min at 2000rpm at 4°C. Cells were fixed with PBS/EDTA 5mM/paraformaldehyde 2% (PFA) for 15min at room temperature. PFA was then rinsed twice by with PBS and removed by centrifugation as described previously, and cells were resuspended by with PBS/EDTA 5mM before quantification of DiI and A680-associated fluorescence with Attune™ NxT flow cytometer equipped with Attune™ Cytometric software v5.2.0.

### Transfection experiments

Neuro-2a cells were plated in a 96-well plate at 4 000 cells/well one day before the experiment. According to DharmaFECT general manufacturer’s protocol (Horizon Discovery), cells were transfected with a mix containing the DharmaFECT 2 and 30nM of tested compound. This mix was incubated on cells in DMEM supplemented with 10% fetal calf serum during 24h at 37°C. At the end of the incubation period, Neuro-2a cells were washed once with D-PBS before RNA extraction.

### Uptake and free uptake gene-silencing experiments

Neuro-2a cells were plated in a 48-well plate at 13 000 cells/well the day before the test. All siRNA and conjugates were incubated during 3 days at 37°C with 1μM of conjugate prepared in DMEM supplemented with 1% fetal calf serum. At the end of the incubation period, cells were washed once with D-PBS before RNA extraction.

### Animal handling

Wild-type male C57Bl/6JRj mice (Janvier Labs, Le Genest-Saint-Isle, France, EU) aged 10-12 weeks old were used. All animal studies were approved by the ethics committee for animal experimentation (CEEA-N°14) and approved by French ministry of agriculture (MRC). During all the experiments, animals were housed per group consisting of 4 animals, on a 12 h light /12 h dark cycle, with food and water access *ad libitum*. Intravenous (i.v., lateral tail vein) administrations (15 mg/kg) were performed on conscious restrained mice. Following a 7-day observational period, animals were euthanized by an intraperitoneal overdose of Ketamin-Xylasine mixture and tissue samples were collected after extensive blood wash-out by intracardiac perfusion (Left ventricle, Flow rate 8ml/min) of a saline solution (0.9%) and immediately stored at −20°C in 10 vol NucleoProtect RNA stabilization reagent (Macherey-Nagel) before further RNA extraction and target mRNA quantification.

### RNA extraction and cDNA synthesis

For *in vitro* free uptake experiments the total RNA was extracted using the NucleoSpin RNA XS (Macherey Nagel) according to the manufacturer’s recommendations. For transfection experiments the SuperScript™ IV CellsDirect™ cDNA Synthesis kit (Invitrogen) was used according to manufacturer’s recommendations. For *in vivo* experiments, mouse organs were crushed in 2mL tubes pre-filled with ceramic mixture in Precellys^®^Cryolys^®^ Evolution (Bertin), with a QIAZOL lysis buffer (Qiagen). A volume of 150μL of chloroform (SIGMA) was added to 750μL of tissue homogenate and a phenol/chloroform separation was performed by centrifugation for 15min at 6000G at 4°C. Aqueous phases were recovered and RNA extraction was performed with the RNeasy 96 QIAcube HT Kit (Qiagen) according to the manufacturer’s recommendations, in a QIACUBE HT instrument (Qiagen). Quality and quantity of total RNA was determined with DNF-471 RNA Kit −15 nt (Agilent) in the Fragment Analyzer 5300 (Agilent). For reverse transcription (RT), 500ng of total RNA were used (except for cells treated with SuperScript™ IV CellsDirect™ cDNA Synthesis), and cDNA synthetis was performed using the High-Capacity RNA-to-cDNA™ kit (Invitrogen) according to the manufacturer’s recommendations.

### Real-time quantitative PCR

The real-time quantitative PCR (qPCR) assays were performed using the CFX96 Touch Real-Time PCR Detection System (Bio-Rad). Amplifications were carried out in a 10 μL final reaction solution containing 12.5ng of cDNA, 1X of TaqMan™ Fast Universal PCR Master Mix (2X), no AmpErase™ UNG (Invitrogen), 1X of TaqMan^®^ Gene Expression Assay Mix (Life Technologies) and RNase free water, according to the manufacturer’s recommendations. The following primers were used: Mm01344233_g1 SOD1(mouse), Mm02526700_g1 RpL13 (mouse), Mm01352366_m1 SDHA (mouse), Mm00457191_m1 PSMC4 (mouse). RpL13, SDHA or PSMC4 served as internal controls for sample normalization and the comparative cycle threshold method (2^−ΔΔCt^) was used for data quantification. Finally, gene expression ratios (compared to control samples) were determined.

### Data analysis

Statistical comparison of the knock-down effect of tested molecules in both *in vitro* free uptake and *in vivo* experiments was performed using a one-way ANOVA followed by a Dunnett’s multiple comparisons test.

*In vivo* experiments were performed blind from treatment administration until data analysis and freezing (all formulations and tissue samples were coded).

## Notes

### Competing Interest Statement

The authors have declared no competing interest.

