## Supplementary figures and images for "LDLR-mediated targeting and productive uptake of siRNA-peptide ligand conjugates *in vitro* and *in vivo*"

### Supplemental Table 1

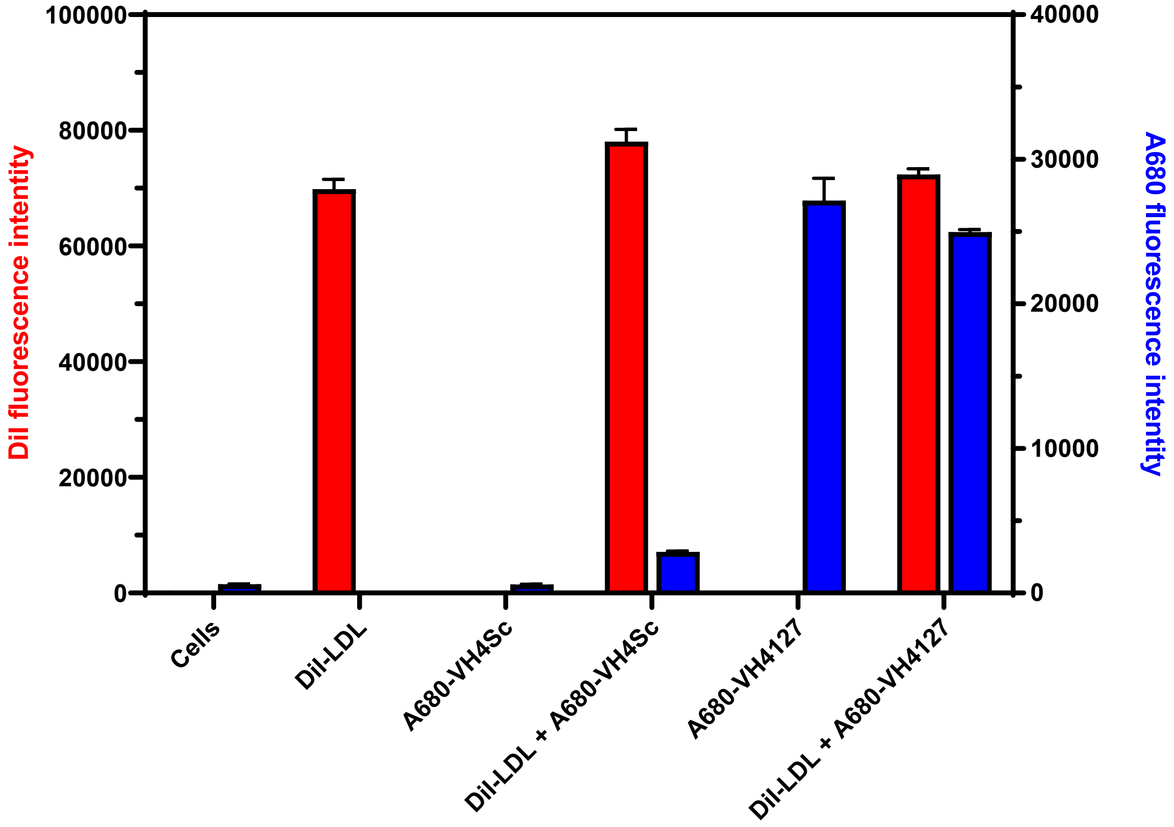


**Supplemental Figure: in vitro validation of LDLR-expressing Neuro-2a cell line.**
